# Building an open infrastructure for molecular neuroimaging: standards and tools from the OpenNeuroPET initiative

**DOI:** 10.64898/2026.05.06.722876

**Authors:** Melanie Ganz, Martin Norgaard, Cyril Pernet, Granville J. Matheson, Anthony Galassi, Eric Ceballos, Murat Bilgel, Cyrus Eierud, Gabriel Gonzalez-Escamilla, Paul Wighton, Christopher J. Markiewicz, Nell Hardcastle, Joshua Buckholtz, Ross Blair, Douglas N. Greve, Adam G. Thomas, Russell A. Poldrack, Vince Calhoun, Robert B. Innis, Gitte M. Knudsen

## Abstract

Molecular neuroimaging with positron emission tomography (PET) and single-photon emission computed tomography (SPECT) enables quantification of specific molecular targets in the living brain. Despite its scientific impact, molecular neuroimaging research has historically faced challenges due to high costs, small sample sizes, laboratory-specific analysis pipelines, and limited large-scale data sharing. These factors have hindered reproducibility and the broader reuse of valuable PET datasets.

The OpenNeuroPET initiative was established to address these barriers by developing standards, infrastructure, and open-source tools currently focused on PET. Through collaborations across Europe and North America, OpenNeuroPET has supported the PET extension of the Brain Imaging Data Structure (PET-BIDS), providing a standardized framework for PET datasets and metadata. Building on PET-BIDS, tools such as PET2BIDS, ezBIDS, and BIDSCoin facilitate data conversion and curation. In parallel, OpenNeuro now hosts PET-BIDS datasets for open sharing, while complementary platforms such as PublicnEUro provide controlled-access pathways designed to support compliance with the European Union General Data Protection Regulation (GDPR). Emerging open-source workflows further support automated, reproducible PET analysis, promoting harmonization across centers.

Together, these developments mark an important step toward an open molecular neuroimaging community in which datasets, software, and workflows can be transparently shared, reused, and scaled for collaborative research.

## Introduction

Molecular neuroimaging with positron emission tomography (PET) and single-photon emission computed tomography (SPECT) enables in vivo measurement and quantification of brain metabolism, neurotransmitter systems, neuroinflammation, synaptic density, protein aggregation, and other molecular targets and processes. These methods have broad applications spanning fundamental neuroscience, clinical research, and drug development. However, they are also extremely costly, require specialized infrastructure and personnel, and involve exposing participants to ionizing radiation. These factors impose practical limits on sample sizes, while laboratory-specific data formats and analysis pipelines further hinder reuse and reproducibility. Together, these constraints create both scientific and ethical imperatives for making optimal use of collected data. This calls for the adoption of transparent and reproducible analysis practices, together with a move towards multi-site collaboration and data sharing.

At the same time, as neuroimaging research in general has advanced over the past two decades, characterized by larger datasets, multi-site collaborations, and increasingly sophisticated computational analyses, it has encountered substantial challenges related to both practical aspects surrounding these collaborations and how best to harmonise sharing and processing of research data, together with the inherent analytical flexibility of complex datasets and models.^1,2^ These challenges can directly compromise the integrity of quantitative measurements, making it crucial for researchers to implement open, transparent, and robust data management practices.^1^ Given the high costs and technical demands associated with individual PET scans—often amounting to several thousands of dollars per scan and requiring specialized infrastructure—the need for transparency and efficiency in data usage becomes paramount, thus also calling for responsible data sharing. By improving data organization and analysis reproducibility, researchers can enhance the reliability of their findings. This ultimately ensures that advances in molecular neuroimaging contribute meaningfully to translational neuroscience and drug development, benefitting society by fostering improved health outcomes.

Earlier efforts to standardize PET data analysis and reporting laid important groundwork for current developments. These include early methodological frameworks for tracer quantification and cross-center harmonization, as well as attempts to improve reporting consistency and data comparability across laboratories^3,4^. However, these efforts primarily focused on methodological standardization and quantitative analysis rather than on infrastructure for organizing and responsibly sharing complete datasets. In particular, they were not centered on challenges related to data sharing and privacy, and predated the development of the Findable, Accessible, Interoperable, and Reusable (FAIR) principles, which provide a broader framework for effective research data management and stewardship^5^.

In response to these challenges, the neuroimaging community has developed several open standards and infrastructures designed to facilitate reproducible research. The Brain Imaging Data Structure (BIDS) provides a community-driven specification for organizing and describing neuroimaging datasets,^6^ while repositories such as OpenNeuro allow researchers to share standardized BIDS datasets.^7^ In parallel, open-source software frameworks developed by the neuroimaging community have introduced robust, containerized preprocessing pipelines (BIDS Applications) that enable transparent and reproducible neuroimaging analyses.^8^

Until recently, these developments have focused primarily on magnetic resonance imaging (MRI) data, and comparable infrastructure for molecular imaging remained limited. Because molecular imaging studies often rely on laboratory-specific data formats, heterogeneous analysis pipelines, and limited opportunities for large-scale data sharing, addressing these limitations requires a coordinated community effort to extend existing standards and infrastructure to support molecular imaging data. Although many of the principles and tools described in this work apply to both PET and SPECT, the standards and tools reviewed are currently focused on PET, while SPECT remains a future modality-specific extension.

### The OpenNeuroPET initiative: goals and scope

OpenNeuroPET is an international collaboration of research groups across Europe and North America that aims to address the challenges related to data standardization and sharing and accelerate the adoption of open, reproducible practices in molecular neuroimaging. The initiative was launched in 2020 through coordinated funding from the Novo Nordisk Foundation in Europe and the National Institutes of Health (NIH) BRAIN Initiative in the United States.

The core goals of OpenNeuroPET are to:

1. establish community standards for organizing and sharing PET data through PET-BIDS, while supporting future modality-specific extensions for SPECT;
2. develop software to facilitate research groups in the implementation of these standards, including preparation of BIDS-compatible data from raw PET data;
3. extend existing neuroimaging repositories, including OpenNeuro, to support sharing of standardized PET datasets, as well as provide additional General Data Protection Regulation (GDPR)-compatible data sharing frameworks for neuroimaging data;
4. develop open-source software tools that enable robust and reproducible PET data processing and analysis, including training sessions and production of educational material; and
5. enable development and sharing of molecular imaging brain atlases and related derivative resources based on standardized, reusable PET data.

The NIH-funded component of the project primarily focuses on integrating PET datasets into the U.S.-based OpenNeuro platform and enabling interoperability with the National Institute of Mental Health’s National Data Archive (NIMH NDA). In parallel, the Novo Nordisk Foundation–funded component focuses on developing GDPR-compatible infrastructure for PET data sharing in Europe (PublicnEUro). Both efforts have contributed to the development of the BIDS standard, data analysis tools, and training materials designed to support open molecular imaging research.

Importantly, OpenNeuroPET has evolved into a broader community effort that extends beyond the initially funded projects. Researchers and developers from multiple institutions have helped develop standards, open datasets, and analysis software, reflecting the strong collaborative culture within the molecular neuroimaging community.

The following sections describe the key components of this infrastructure, including the development of data standards, data-sharing infrastructure, and open-source curation and analysis tools that, together with training and educational material, support reproducible molecular neuroimaging research.

### Standardization of PET data and metadata

Efforts to standardize molecular neuroimaging data began within the Neuroreceptor Mapping (NRM) community, which established consensus terminology for reporting kinetic modeling outcomes as early as 2007.^9^ However, standardized reporting of the full experimental setup of PET and SPECT studies—including tracer synthesis, acquisition protocols, and blood measurements—emerged much later. These reporting standards were developed through a multi-year community effort initiated at the NRM meeting in 2016 and coordinated by Dr. Robert B. Innis and Prof. Gitte Moos Knudsen. The process involved contributions from multiple sub-disciplines within molecular imaging, including radiochemists, physicists, neuroscientists, clinicians, and nuclear medicine technologists. The resulting consensus guidelines defined recommended practices for reporting PET neuroimaging experiments in both publications and data archives.^10^

PET imaging experiments often involve multiple data streams beyond the reconstructed images acquired from the scanner, and multiple types of expertise. The procedures include radiochemistry with subsequent quality control, radiotracer injection by technicians or physicians, PET data acquisition, and in some cases, blood sampling, where the samples are processed, counted for radioactivity, and analyzed for radiometabolites by specialists. The PET scanner also needs to be cross-calibrated across the gamma counter and the automatic blood sampling device. Bringing together data from these multiple sources requires a highly controlled workflow to avoid mistakes. As a result, PET datasets are inherently multimodal, combining imaging data with biochemical and physiological measurements that vary in scope and complexity depending on the tracer and acquisition choices. In addition, imaging data formats vary across scanners. While Digital Imaging and Communications in Medicine (DICOM)^11^ is the international clinical standard for storing and exchanging medical images and is widely used for magnetic resonance (MR) and computed tomography (CT) scanners, some PET scanners, particularly older systems such as the Siemens Biograph High Resolution Research Tomograph (HRRT), often rely on proprietary formats (e.g., ECAT) for reconstruction and storage, even though DICOM export is typically supported.

Within the MR imaging community, similar challenges related to heterogeneous data formats and incomplete metadata led to the development of BIDS, which was introduced in 2015 and formally published in 2016 as a community-driven standard for organizing and describing neuroimaging datasets.^6^ The standard simplifies data organization, captures essential experimental metadata, and removes participant-identifying information present in raw DICOM headers, thereby facilitating data sharing and reproducible analysis. Following its initial release for MRI data, the BIDS specification was rapidly extended to additional neuroimaging modalities, including magnetoencephalography (MEG)^12^ and electroencephalography (EEG).^13^

Building upon the previously established guidelines paper for the content and format of PET brain data in publications and archives,^10^ the molecular imaging community developed an extension of BIDS for PET data (PET-BIDS).^11^ PET-BIDS defines standardized directory structures, file naming conventions, and metadata specifications for PET imaging data as well as associated experimental measurements such as injected dose, tracer information, and blood radioactivity and metabolite data. PET-BIDS documents the injected mass and molar activity together with their units and the time of molar-activity measurement. ^14^ These are important to record, because excessive injected mass may produce non-negligible target occupancy and bias quantitative estimates. Although tracer administration is generally intended to occupy a negligible proportion of target sites (often defined as less than approximately 5–10%) the appropriate mass limit is tracer- and target-specific. ^9^ Within this framework, harmonization across centers should not imply a universal blood-sampling schedule, as these are tracer-specific. Rather, PET-BIDS provides a common representation of sampling times, metabolite-analysis method, parent fraction, plasma radioactivity, and, where measured, plasma free fraction and its measurement method should also be reported.^10,14^ The PET-BIDS specification was incorporated into the official BIDS standard in 2021 and subsequently described in a dedicated publication in 2022.^14^ Importantly, within the BIDS framework, the term raw data refers to reconstructed imaging volumes and associated experimental measurements rather than scanner-level photon count (e.g., list-mode) data.^14,15^

The introduction of PET-BIDS established a standardized representation of molecular imaging experiments, enabling PET datasets to be organized, shared, and processed using common tools and infrastructures. This development provided a crucial foundation for the expansion of PET data sharing platforms and reproducible analysis pipelines, which are described in subsequent sections.

### Converting PET data into PET-BIDS

Although the PET-BIDS specification provides a standardized structure for organizing molecular imaging datasets, converting existing laboratory datasets into this format remains a practical challenge. PET studies are often stored in laboratory-or even researcher-specific directory structures and include multiple heterogeneous data streams, such as reconstructed PET images, tracer injection metadata, and blood measurements. Preparing these datasets for standardized storage and sharing, therefore, requires substantial effort to extract, harmonize, and document metadata.

In the early stages of PET-BIDS adoption, data conversion was typically performed using the widely used DICOM-to-NIfTI conversion tool dcm2niix.^16^ But due to the missing file-naming capability of dcm2niix, researchers commonly relied on additional custom MATLAB or Python scripts to rename and organize files and populate the additional metadata fields required by the PET-BIDS specification. To streamline this process, the OpenNeuroPET initiative and the broader molecular imaging community developed PET2BIDS, an open-source library specifically designed to convert PET experiments into PET-BIDS-compliant datasets.^17^ PET2BIDS builds upon the robust image conversion capabilities of dcm2niix while providing dedicated tools for structuring PET imaging data, incorporating tracer information, and integrating blood measurements and other experimental metadata. By formalizing common conversion steps into reusable software components, PET2BIDS significantly reduced the technical barrier for adopting PET-BIDS.

To investigate real-world PET2BIDS usage, we started collecting anonymized telemetry data in the fall of 2024. As of April 2026, this telemetry data comprises >60,000 execution events (i.e. conversions of PET data to PET-BIDS) across at least seven countries and ∼20 sites. Usage was heterogeneous over time but remained consistently sustained throughout the observation period, with a mean of ∼732 weekly executions (standard deviation ∼1551), indicating active adoption across multiple research centers.

In parallel, the broader BIDS ecosystem developed graphical user interface (GUI)–based tools that make dataset curation accessible to researchers without extensive programming experience. Platforms such as ezBIDS^18^ and BIDScoin^19^ provide interactive workflows that guide users through the process of mapping raw imaging data into BIDS-compliant datasets.^18,19^ Through integration with PET2BIDS and related libraries, these tools now support PET datasets alongside other neuroimaging modalities, enabling researchers to convert and validate PET-BIDS datasets using web-based or graphical interfaces rather than command-line scripts.

Despite these advances, challenges remain because molecular imaging experiments require physical and biological measurements acquired outside the scanner environment, such as injected radioactivity and blood sampling. These data are often recorded manually or stored in separate laboratory systems, complicating their integration into standardized datasets. Emerging tools aim to address this limitation by capturing or structuring experimental metadata more systematically. For example, eCOBIDAS provides an interactive checklist framework for reporting methods,^20^ and the broader ReproSchema ecosystem supports standardized, schema-based metadata collection that can be integrated with acquisition and curation workflows.^21^ Operating on locally hosted databases compatible with hospital IT infrastructure, such systems can later be integrated with automated conversion pipelines.

Together, these developments illustrate the transition from ad hoc conversion scripts towards increasingly integrated and automated workflows for PET dataset organization. Continued improvements in conversion tools and metadata handling will further facilitate the adoption of PET-BIDS, supporting scalable and reproducible molecular neuroimaging research.

### Sharing PET data: open and controlled-access pathways

A central contribution of the OpenNeuroPET initiative has been the development and extension of platforms for sharing molecular neuroimaging data. Historically, PET data sharing was largely determined by laboratory policies and was often performed only when required by funding agencies. Early efforts nevertheless demonstrated the scientific value of shared molecular imaging datasets. For example, the Neurobiology Research Unit at Copenhagen University Hospital established the Center for Integrated Molecular Brain Imaging (CIMBI) database,^22^ which hosts a large number of PET data (>1000 participants). While this resource enables important collaborations, access to the data is limited: querying the database and navigating its metadata requires dedicated data management support and typically involves direct interaction with individual study investigators.

Finalizing standardized data organization through PET-BIDS enabled the development of more scalable and transparent sharing platforms. In 2022, PET support was formally added to the OpenNeuro repository,^7,14^ creating a dedicated portal for PET datasets. Since then, PET datasets representing more than 1,700 participants across more than 30 datasets have been shared on the platform. These datasets span 24 distinct radiotracers and originate from 19 imaging sites across nine countries (the United States, Canada, Belgium, Germany, China, Australia, the United Kingdom, Switzerland and Denmark), reflecting the growing geographic and methodological diversity of openly shared PET data (Table 1). The collection also includes datasets from multiple species, including humans, non-human primates, rats, and mice. Some datasets were deposited in OpenNeuro before the PET portal was introduced, and the rate of data deposition has increased markedly following its launch.

**Table 1.** Overview of 33 PET datasets currently shared on OpenNeuro that conform to the PET-BIDS standard. For each dataset, the table lists the OpenNeuro dataset identifier (DS00XXXX), PET scanner model, radiotracer(s) used, contributing site and country, number of subjects (N), and species.

|  |  |  |  |  |  |
| --- | --- | --- | --- | --- | --- |
| DS00XXXX | Scanner | Radiotracer | Site/Country | N | Species |
| 7907 | Siemens Biograph mMR | [<sup>11</sup>C]PBR28 | Martinos Center/United States | 147 | Human |
| 7768 | Siemens Biograph Vision Quadra long-axial-field-of-view PET/CT | [<sup>18</sup>F]FDG | University Hospital of Psychiatry Zurich, Switzerland | 14 | Human |
| 7561 | Siemens Biograph Horizon | [<sup>11</sup>C]UCB-J | University of Wisconsin-Madison/United States | 20 | Human |
| 7135 | Siemens BrainPET insert | [<sup>11</sup>C]Martinostat | Martinos Center/United States | 25 | Human |
| 6917 | UIH uNeuroX & Siemens HRRT | [<sup>18</sup>F]FDG; [<sup>18</sup>F]SynVesT-1; [<sup>18</sup>F]FPEB; [<sup>18</sup>F]Flubatine; [<sup>11</sup>C]PHNO; [<sup>18</sup>F]FE-PE2I; [<sup>11</sup>C]DASB | Yale PET Center/United States | 7 | Human |
| 6756 | Siemens HRRT | [<sup>18</sup>F]MK6240 | McConnell Brain Imaging Centre/Canada | 33 | Human |
| 6691 | Siemens Inveon MPET | [<sup>18</sup>F]CHDI-385 | University of Antwerp/Belgium | 88 | Mice |
| 6613 | Siemens Inveon MPET | [<sup>11</sup>C]CHDI-009R | University of Antwerp/Belgium | 95 | Mice |
| 6218 | Siemens Inveon DPET | [<sup>18</sup>F]SynVesT-1 | University of Antwerp/Belgium | 138 | Mice |
| 6148 | Siemens Biograph64 mCT | [<sup>18</sup>F]FDG<br>[<sup>18</sup>F]MK6240<br>[<sup>18</sup>F]florbetaben | Cornell University/United States | 266 | Human |
| 5895 | Molecubes NV Beta-CUBE | [<sup>18</sup>F]FE-PE2I | KU Leuven/Belgium | 25 | Rat |
| 5698 | Siemens Biograph mCT | [<sup>18</sup>F]PF-06445974 | MIB NIMH/United States | 25 | Human |
| 5590 | Siemens PETmCT | [<sup>18</sup>F]SF12051 | MIB NIMH/United States | 7 | Human |
| 5605 | Mediso LFER | [<sup>11</sup>C]MC1 | MIB NIMH/United States | 18 | Rat |
| 5483 | Siemens PETmCT | [<sup>11</sup>C]PS13 | MIB NIMH and Stonybrook/United States | 33 | Human |
| 5093 | Siemens HRRT | [<sup>11</sup>C]PBR28 | University of British Columbia/Canada | 11 | Non-human primate |
| 4869 | Siemens Biograph mCT | [<sup>11</sup>C]MC1 | MIB NIMH/United States | 27 | Human |
| 4868 | Siemens Biograph mCT | [<sup>11</sup>C]PS13 | MIB NIMH/United States | 10 | Human |
| 4856 | Siemens 962 | [<sup>18</sup>F]AV45<br>[<sup>18</sup>F]AV1451 | University of Texas Southwestern Medical Center/United States | 464 | Human |
| 4733 | Siemens HRRT | L-[1-<sup>11</sup>C]Leucine | NIMH/United States | 18 | Human |
| 4731 | Siemens HRRT | L-[1-<sup>11</sup>C]Leucine | NIMH/United States | 35 | Human |
| 4730 | Siemens HRRT | L-[1-<sup>11</sup>C]Leucine | NIMH/United States | 10 | Human |
| 4654 | Siemens HRRT | L-[1-<sup>11</sup>C]Leucine | NIMH/United States | 28 | Human |
| 4513 | Siemens Biograph mMR | [<sup>18</sup>F]FDG | TUM/Germany; | 30 | Human |
| 4230 | Siemens Biograph64 mCT | [<sup>11</sup>C]PS13 | MIB NIMH/United States | 16 | Human |
| 4054 | Siemens Biograph mMR | [<sup>18</sup>F]FDG | Shanghai Tech University/China | 42 | Human |
| 3397 | Siemens Biograph mMR | [<sup>18</sup>F]FDG | Monash University/Australia | 15 | Human |
| 3382 | Siemens Biograph mMR | [<sup>18</sup>F]FDG | Monash University/Australia | 10 | Human |
| 2898 | Siemens Biograph mMR | [<sup>18</sup>F]FDG | Monash University/Australia | 27 | Human |
| 2041 | GE Discovery STE | [<sup>18</sup>F]Fallypride | Duke University/United States | 25 | Human |
| 1705 | Siemens Biograph mMR | [<sup>11</sup>C]LondonPride | KCL/United Kingdom | 5 | Human |
| 1421 | Siemens HRRT | [<sup>11</sup>C]SB207145 | Neurobiology Research Unit/Denmark | 1 | Human |
| 1420 | Siemens HRRT | [<sup>11</sup>C]DASB | Neurobiology Research Unit/Denmark | 2 | Human |

Figure 1 illustrates the cumulative growth of publicly available PET participants on OpenNeuro between 2018 and 2026 (June).

**Figure 1:**
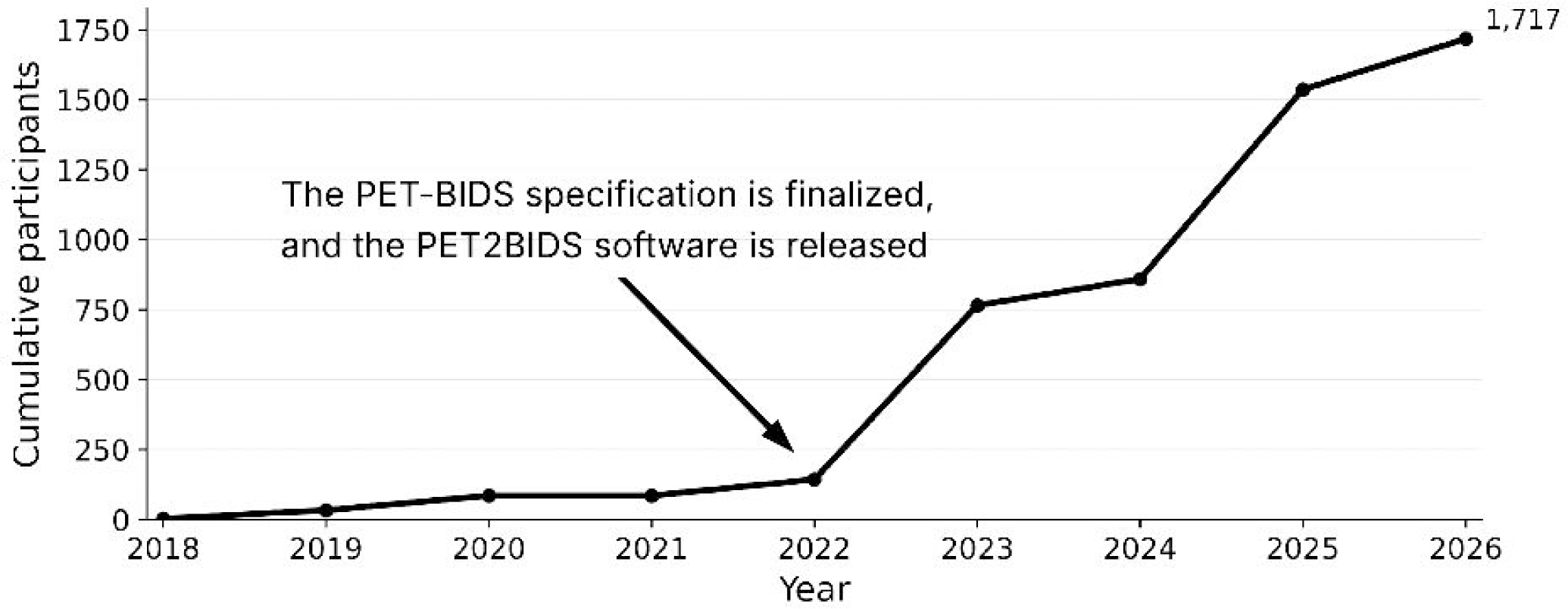
Cumulative number of participants per year represented in PET datasets publicly available through OpenNeuro between July 2018 and June 2026. In total, the figure includes 1,717 subjects from 33 datasets.

Despite the advantages of fully open data sharing, regulatory requirements— particularly within the European Union—often limit the possibility of releasing neuroimaging datasets under unrestricted licenses such as the Creative Commons Zero (CC0) license used by OpenNeuro. To address this limitation, a complementary GDPR-compliant platform, PublicnEUro, was developed as part of the European component of the OpenNeuroPET initiative.^23,24^ PublicnEUro received its first dataset deposits in 2025 and has since expanded to host 17 datasets comprising more than 500 participants (Table 2).

**Table 2.** Overview of 17 PET datasets currently shared on PublicnEuro that conform to the PET-BIDS standard. For each dataset, the table lists the PublicnEUro dataset identifier (PNXXXX), PET scanner model, radiotracer(s) used, contributing site and country, number of subjects (N), and species. One peculiar dataset is PN000001, an open-access repository of source phantom data curated to BIDS, allowing software testing for data conversion.

| PNID | Scanner | Radiotracer | Site/Country | N | Species |
| --- | --- | --- | --- | --- | --- |
| PN000001 | Siemens, HRRT, Biograph One, Biograph MRM, Vision quadra and Brain Trio<br>Canon Cartesion, GE Advance, Discovery, and Signa, Phillips Ingenuity and Vereo | [<sup>18</sup>F]FDG | Denmark, USA, Germany, Netherlands | 18 | Phantom data |
| PN000002 | Siemens HRRT | 5-HT1BR | NRU/Denmark | 50 | Human |
| PN000005 | Siemens HRRT | [<sup>11</sup>C]UCB-J | Leuven/Belgium | 12 | Human |
| PN000006 | Siemens HRRT | [<sup>11</sup>C]UCB-J | Leuven/Belgium | 11 | Human |
| PN000007 | Siemens HRRT | [<sup>11</sup>C]UCB-J | Leuven/Belgium | 40 | Human |
| PN000008 | Siemens HRRT | [<sup>11</sup>C]UCB-J | Leuven/Belgium | 20 | Human |
| PN000010 | Siemens HRRT | [<sup>11</sup>C]UCB-J | Leuven/Belgium | 15 | Human |
| PN000012 | Siemens HRRT | [<sup>11</sup>C]UCB-J | NRU/Denmark | 41 | Human |
| PN000012 | Siemens HRRT | [<sup>11</sup>C]UCB-J<br>[<sup>18</sup>F]FDG | Aarhus/Denmark | 63 | Human |
| PNC00002 | Siemens Quadra | [<sup>18</sup>F]FDG | Rigshopitalet/Denmark | 80 | Human |
| PN000017 | Siemens HRRT | [<sup>11</sup>C]DASB | NRU/Denmark | 53 | Human |
| PN000018 | Siemens HRRT | [<sup>11</sup>C]DASB | NRU/Denmark | 63 | Human |
| PN000019 | Siemens HRRT | [<sup>11</sup>C]DASB | NRU/Denmark | 24 | Human |
| PN000020 | Siemens HRRT | [<sup>11</sup>C]DASB | NRU/Denmark | 46 | Human |
| PN000021 | Siemens HRRT | [<sup>11</sup>C]DASB | NRU/Denmark | 33 | Human |
| PN000022 | Siemens HRRT | [<sup>11</sup>C]DASB | NRU/Denmark | 21 | Human |
| PN000023 | Siemens HRRT | [<sup>11</sup>C]DASB | NRU/Denmark | 10 | Human |

Unlike OpenNeuro, which distributes datasets directly under open licenses, PublicnEUro operates as an intermediary infrastructure between data owners and data users. The platform provides secure storage coupled with data owner-centric transfer mechanisms.

Access to each dataset is governed by the data owners, allowing each institution to fulfil its legal requirements while automating the process, thus increasing data accessibility. At the same time, PublicnEUro exposes standardized metadata that can be publicly browsed, thereby improving findability.

Looking forward, an important next step will be enabling unified discovery across multiple neuroimaging repositories. Emerging tools such as the Neurobagel ecosystem^25^ support distributed metadata harmonization and federated searches across independent databases. This approach will allow researchers to identify relevant datasets across platforms, including OpenNeuro, PublicnEUro, and the EBRAINS knowledge graph, based on characteristics such as radiotracers, participant demographics, or disease conditions, moving towards a more interoperable global ecosystem for molecular neuroimaging data sharing.

### Reproducible processing of PET data: from preprocessing to kinetic modeling

The analysis of PET data has historically relied on heterogeneous workflows composed of multiple independent software tools,^26–28^ including widely used neuroimaging packages such as Automated Image Registration (AIR), Statistical Parametric Mapping (SPM), FMRIB Software Library (FSL), and FreeSurfer,^29–32^ often combined with specialized proprietary software for quantification, such as PMOD. While these tools enabled major scientific advances in molecular neuroimaging, the lack of standardized pipelines and reproducible computational environments has made analyses difficult to reproduce across laboratories. The impact of analytical flexibility in PET processing has been highlighted by several community initiatives, including the NRM Grand Challenge, which demonstrated substantial variability in quantitative outcomes depending on preprocessing choices made when analysing the same dataset.^33,34^

### Preprocessing of PET data

Early PET preprocessing workflows were typically assembled from tools originally developed for structural and functional MRI analysis. Packages such as AIR, SPM, FSL, and FreeSurfer were commonly used for motion correction, spatial normalization, PET– MRI co-registration, and anatomical segmentation, while later extensions such as PETSurfer^35,36^ introduced PET-specific processing within existing neuroimaging environments.^37^ In parallel, proprietary platforms such as PMOD provided integrated environments that simplified preprocessing and quantification through graphical interfaces, but their closed-source implementations and limited automation constrained transparency and reproducibility.

More recently, the PET analysis ecosystem has begun shifting toward standardized and reproducible workflows. This transition was strongly influenced by BIDS and by the emergence of BIDS applications.^8,14^ Tools such as PETPrep (https://petprep.readthedocs.io/en/latest/) and related PET BIDS-applications (e.g., PICNIC^38^) aim to automate common preprocessing steps—including motion correction, PET–MRI co-registration, spatial normalization, region-of-interest definition, and optional partial volume correction—while ensuring consistent data organization and metadata handling.^14,39^ By encoding analysis decisions within version-controlled workflows that can be executed in containerized environments (e.g., Docker (https://www.docker.com/) or Apptainer (https://apptainer.org/)), these pipelines reduce manual intervention and improve reproducibility across datasets and computing infrastructures.

Future developments will likely focus on closer integration between preprocessing pipelines and downstream analysis tools, improved interoperability across software ecosystems, and scalable processing of large multi-site datasets. Additional advances may include automated quality control, standardized derivative outputs, and integration with cloud-based computing platforms.

### Pharmacokinetic modeling of PET data

Quantification of PET data relies on pharmacokinetic modeling to estimate parameters describing tracer binding, distribution, or metabolism from dynamic imaging data and associated measurements such as blood input functions. Historically, kinetic modeling has relied on commercial platforms, laboratory-specific software, and publicly distributed academic tools. PMOD provides a mature, integrated environment with extensive modeling and visualization functionality. However, its proprietary implementation prevents independent source-code inspection, requires licensing, and may complicate exact reproduction within automated, version-controlled workflows. These limitations concern transparency and computational reproducibility rather than the scientific validity of the implemented methods.

Long-standing academic resources include the Turku PET Centre command-line programs (http://www.turkupetcentre.net/petanalysis/index.html), which implement many established kinetic models and related utilities, although interfaces, source availability, licensing, and maintenance vary across individual programs. More recent open and source-available tools increasingly support scripted, version-controlled, and containerized analyses.

Representative PET preprocessing and kinetic-modeling resources are summarized in Supplementary Table S1.

Open-source kinetic modeling tools have gradually emerged over the past decade, although early implementations were often fragmented and lacked standardized input formats. More recently, several tools have begun integrating modeling with standardized data structures and reproducible workflows. Examples include kinfitr,^40^ PETFit,^41^ and modeling functionality within PETSurfer-based workflows.^37^ These tools support visualization of data and model fitting, parametric imaging, and blood input function handling while enabling analyses to be scripted, version-controlled, and incorporated into automated pipelines.

Continued development of open kinetic modeling frameworks and associated training will be essential for establishing fully reproducible PET analysis workflows. Future directions include tighter integration between preprocessing and modeling tools, standardized representations of modeling inputs and outputs within BIDS derivatives, and improved support for large-scale analyses across multi-site datasets. Together, these developments move the field toward a comprehensive open ecosystem for molecular neuroimaging analysis spanning preprocessing, quantitative modeling, and downstream statistical analysis.

### Molecular imaging brain atlases: from shared derivatives to reusable reference resources

An additional goal of the OpenNeuroPET initiative is to support the creation and sharing of molecular imaging brain atlases (MIBAs). In this context, MIBAs are standardized group-level PET or SPECT derivative resources describing the spatial distribution of molecular targets in the brain, including receptors, transporters, enzymes, protein aggregates, and synaptic markers (Figure 2). These atlases may be represented as voxel-wise or surface-based maps, region-wise summaries, or multivariate component maps and serve as reusable resources for visualization, hypothesis generation, benchmarking, and downstream statistical analysis. As openly available datasets increase, the field is transitioning from sharing primary data alone towards distributing curated derivative resources that capture molecular organization across cohorts.^42–44^

**Figure 2:** Examples of Molecular Imaging Brain Atlases (MIBAs) created from publicly available data on OpenNeuro. The first is a COX-2 atlas using the radiotracer [^11^C]MC1 from the dataset ID ds004869 (https://openneuro.org/datasets/ds004869/) on OpenNeuro (left), and the other is a PDE4B atlas using the radiotracer [¹⁸F]PF-06445974 from the dataset ID ds005698 (https://openneuro.org/datasets/ds005698/) on OpenNeuro.

Within the BIDS ecosystem, derivative datasets can be broadly divided into subject-level outputs and group-level resources, with MIBAs belonging to the latter. Although PET atlases have been developed before, their dissemination has historically been fragmented, often limited to laboratory websites or supplementary materials without standardized metadata or versioning. Recent extensions of the BIDS specification introduce formal support for atlas derivatives across modalities, enabling standardized representation and sharing of such resources.^45,46^ This provides a foundation for distributing PET atlases through community repositories, linking them to source datasets and processing workflows, and facilitating reuse across studies and software tools. In parallel, specifications for subject-level PET derivatives remain under active development and are expected to mature alongside atlas standards.^14,46^

Several studies highlight the scientific value of MIBAs. Early work produced high-resolution atlases of specific neurotransmitter systems, including multidimensional maps of the human serotonergic system derived from multiple receptor and transporter measurements, including calibration from PET radiotracer-specific measures to actual density.^47,48^ Subsequent studies extended this approach to other targets, such as benzodiazepine receptors^46^ and synaptic density using synaptic vesicle glycoprotein 2A (SV2A) PET.^43^ At a larger scale, data from more than 1,200 healthy individuals have been integrated into a whole-brain atlas of 19 receptors and transporters across nine neurotransmitter systems, demonstrating how MIBAs can be used to relate molecular architecture to macroscale brain organization.^42^ The high number of citations these studies retrieve, typically in the order of 300 citations per year, illustrates that MIBAs are not only descriptive maps but are integrative resources linking PET measurements to structural, functional, and transcriptomic brain organization.^43,47–49^

Within OpenNeuroPET, MIBAs are relevant in two complementary ways. First, the initiative contributes to the technical foundations required to build them reproducibly, including standardized PET-BIDS datasets, harmonized metadata, shared preprocessing workflows, and emerging conventions for derivative outputs. Second, it provides infrastructure for sharing atlas resources themselves. For example, molecular imaging atlas derivatives have already been distributed through OpenNeuro, including a PET atlas of cyclooxygenase-1, illustrating how atlas-style outputs can be shared through community repositories rather than remaining confined to individual laboratories.^47^ This infrastructure can provide a Digital Object identifier (DOI) and supports versioning, documentation of generation workflows, and explicit linkage to source data, thereby improving transparency and reuse.

Methodologically, MIBAs are not limited to regional averages based on anatomical parcellations and voxel-wise maps. Recent approaches derive atlas representations from the covariance structure of molecular imaging data. For example, the NeuroMark-PET framework uses spatially constrained independent component analysis, as implemented in the GIFT software (http://trendscenter.org/software/gift),^51,52^ to construct multivariate atlases with overlapping components that may better capture molecular organization than traditional region-based approaches.^44^

Looking forward, systematic generation of MIBAs may become one of the most important products of openly shared PET data. Standardized workflows could enable the creation of normative reference maps across tracers, targets, age groups, and species, as well as disease-specific and multimodal atlases. When linked to BIDS derivatives and shared through platforms such as OpenNeuro and PublicnEUro, MIBAs can serve as a bridge between primary data sharing and large-scale secondary reuse, extending the OpenNeuroPET vision from standardized datasets and pipelines toward reusable community knowledge resources.

It is important to note that in order to be useful to the community, MIBAs should clearly report the reference population and processing pipeline, including age and sex distributions and whether and how partial-volume correction was applied. Where adequately powered, age-adjusted and sex-adjusted or sex-stratified estimates should be provided. Furthermore, depending on the tracer and research question, additional participant variables such as body mass index, blood pressure or heart rate, hematocrit, smoking status, medication or substance use, and education, should be retained when available and relevant; otherwise, the demographic limitations of the atlas should be stated explicitly.

### Training, documentation, and community building

A core component of the OpenNeuroPET initiative has been community training and openly shared educational material. To support adoption of PET-BIDS, data sharing, and reproducible PET workflows, OpenNeuroPET maintains a public outreach repository containing lectures, slides, workshop materials, and onboarding resources, alongside project webpages that explicitly position training and outreach as part of the initiative’s mission.^53^

Based on the documented event history provided in the outreach repository,^53^ OpenNeuroPET has contributed to 24 training and outreach events between 2021 and 2025, corresponding to an average of 4.8 events per year. Across this period, activities spanned major international conferences, hackathons, data standards meetings, onboarding workshops, and domain-specific PET events, indicating sustained engagement across both broad neuroimaging audiences and more specialized molecular imaging communities.

Several events also reflect recurring engagement with key community venues. For example, OpenNeuroPET-related training or dissemination activities were represented at the Organization for Human Brain Mapping (OHBM) Annual Meeting in 2021, 2022, and 2024, Brainhack Nordic in 2021, 2022, and 2024, NRM in 2021 and 2024, and PET is Wonderful in 2021 and 2025. This repeated presence suggests that community building was not limited to one-off presentations, but rather formed part of a longer-term strategy for developing PET-BIDS literacy, gathering feedback from users, and aligning tooling and standards development with community needs. The outreach materials themselves further show that training covered both conceptual topics, such as PET-BIDS and data sharing, and practical topics, including PETSurfer, PET2BIDS, and workflow-based processing. Another strong example of OpenNeuroPET’s outreach efforts was the Nordic PET-BIDS onboarding and ezBIDS workshop hosted by the Karolinska Institutet in August 2025. The workshop was explicitly designed as hands-on training for PET researchers, covering what BIDS is, how it supports reproducible analysis and data sharing, how to use ezBIDS and PET2BIDS for data curation, how to run PET-specific BIDS applications, and how European researchers can navigate data-sharing options. Importantly, the workshop materials also articulate a community goal: to build a network of PET researchers who can support one another in the transition to BIDS-based workflows. This event was also supported by the International Neuroinformatics Coordinating Facility (INCF), providing an open educational space.^54^

Taken together, these activities highlight that OpenNeuroPET has supported adoption not only by releasing infrastructure and datasets, but also by investing in the social and educational work required for community support of the standard and critical assessment of modeling approaches. In this sense, the initiative’s training program functions as an important bridge between data specification development, software development, and practical use by the PET community.

## Discussion and Outlook

The developments described above illustrate a rapid transition in molecular neuroimaging towards standardized data organization, open data sharing, and reproducible computational workflows. Through the coordinated efforts of the OpenNeuroPET initiative and the broader neuroimaging community, key components of this infrastructure—data standards, repositories, analysis tools, and training—have matured substantially over the past several years. At the same time, several challenges remain before a fully reproducible end-to-end molecular imaging workflow can be realized (Figure 3).

**Figure 3:** Overview of OpenNeuroPET efforts, providing tools to move data from scanner to outcome using BIDS data, ranging from conversion tools, data sharing options, and preprocessing/quantification tools.

One frequently raised concern regarding open data sharing is the risk of data misuse, including the possibility of “scooping” or misinterpreting shared datasets. Similar concerns have been discussed across many areas of science and are not unique to molecular imaging. Importantly, scientific misconduct and analytical errors have always existed, regardless of whether data are shared. Open and reproducible research practices provide mechanisms that can mitigate such risks rather than amplify them. Transparent data organization, open analysis code, and reproducible computational workflows allow analyses to be scrutinized, replicated, and improved by the broader community. Increasingly, funding agencies and journals are also recognizing and incentivizing such practices, reinforcing the view that openness and reproducibility strengthen scientific reliability rather than threaten it.

Another important challenge concerns the computational environments used to analyze PET data. Although containerization technologies such as Docker and Singularity have made it possible to distribute reproducible preprocessing pipelines, many published studies still lack fully documented computational environments for downstream analyses and statistical modeling. Therefore, ensuring long-term reproducibility requires not only standardized data formats but also well-defined computational environments that capture software dependencies, analysis scripts, and workflow configurations. Continued community efforts will be necessary to promote the adoption of containerized workflows and workflow management systems throughout the full PET analysis lifecycle.

The sustainability of open molecular imaging infrastructure also depends on recognizing the resources required to maintain it. Data storage, curation, and software development represent substantial ongoing investments that are often underrepresented in traditional grant budgets, which historically focus primarily on data acquisition. As the field moves towards large shared datasets and collaborative analysis infrastructures, funding mechanisms and institutional policies will need to better support long-term data stewardship and the development of open scientific software.

From a methodological perspective, molecular imaging analyses are also evolving. New PET data analysis models are being introduced, and they need to be properly validated and implemented. Historically, PET studies often relied on relatively simplified statistical approaches, typically focusing on region-of-interest analyses with modest sample sizes and limited correction for multiple comparisons. More recent work has highlighted the risks of inflated false positive rates and the broader implications of analytical flexibility across preprocessing, modeling, and statistical analysis choices.^33,55^ Addressing these issues will likely require more systematic evaluation of analysis pipelines, including multiverse-style approaches that explicitly quantify the impact of alternative analytic decisions. In the long run, integrating preprocessing, kinetic modeling, and statistical inference into coherent and reproducible analysis pipelines may help reduce analytical variability and improve the robustness of PET findings.

While reconstructed imaging data are now standardized through PET-BIDS, scanner-level raw acquisition data, such as PET list-mode data or MR k-space data, remain outside the scope of the current BIDS specification. Similar challenges have been addressed in magnetic resonance imaging through initiatives such as the International Society for Magnetic Resonance in Medicine (ISMRM) Raw Data format (ISMRM-RD),^56^ and discussions are underway within the molecular imaging community to define comparable standards for pre-reconstruction PET data formats.^57^ Standardizing raw acquisition data would represent an important step towards fully transparent molecular imaging workflows, enabling reproducibility not only of downstream analyses but also of reconstruction procedures.

Finally, the growing availability of openly shared PET datasets is beginning to enable new forms of scientific analysis. Public PET datasets organized according to PET-BIDS can now be processed using standardized pipelines and analyzed collectively across studies. Such datasets provide opportunities for methodological benchmarking, cross-tracer comparisons, and the development of normative reference models of molecular imaging signals across the lifespan. As the number of available datasets increases, these resources will likely support increasingly sophisticated statistical analyses and meta-analytic approaches that were previously difficult to perform in molecular imaging due to limited data availability, as well as the development of more sophisticated ways to more fully harmonise outcomes from data collected at different research centres.

Taken together, the advances described above demonstrate that molecular neuroimaging is entering a new phase characterized by standardized data organization, open infrastructure, and reproducible computational workflows. Continued collaboration across laboratories, funding agencies, and infrastructure developers will be essential for sustaining this progress. The increasing number of PET datasets on OpenNeuro and PublicnEUro illustrate the growing adoption of PET-BIDS and enthusiasm for open data sharing in the community, highlighting the impact of the OpenNeuroPET Initiative. If these efforts continue, the coming decade may see the emergence of a fully interoperable infrastructure for molecular imaging research in which data, software, and analyses can be transparently shared, reproduced, and extended across the global scientific community.

## Data availability statement

All data presented in this work is openly available via OpenNeuro, PublicnEUro and GitHub.

