## Supplemental Table 1 for "Building an open infrastructure for molecular neuroimaging: standards and tools from the OpenNeuroPET initiative"

**Supplementary Table S1. Representative historical and current software resources for PET preprocessing, quantitative analysis, and kinetic modelling.**

*Table note. This table is non-exhaustive and is not a ranking of scientific validity, feature completeness, validation quality, or current maintenance. Software descriptions and access conditions were compiled from the cited literature, project pages or repositories, and the overview by Thielemans (2023). Thielemans explicitly described the categorisation as a subjective assessment of open-source criteria rather than an assessment of features or testing. Licences, versions, dependencies, access routes, and project status can change. Publicly available source code, free-to-use software, and open-source software are not equivalent. Current project pages were checked on 13 August 2026 where available.*

| **Tool/resource** | **Period / reference** | **Primary scope** | **Access / licence** | **Reproducibility strengths** | **Caveats / positioning** |
| --- | --- | --- | --- | --- | --- |
| **Early Java PET quantitation environment** | Mikolajczyk et al., 1998; PMID 9785322 | Portable modular medical-image analysis environment with an initial brain PET kinetic-quantitation application. | Published research framework; historical software availability and licence should be checked. | Early emphasis on portability, modularity, and separation from scanner-specific systems. | Historically important; not a contemporary maintained end-to-end workflow. |
| **Turku PET Centre programs** | Long-standing academic suite; Turku PET Centre PET analysis pages | Command-line utilities for TACs, input functions, kinetic models, parametric analysis, and ancillary PET processing. | Publicly distributed; terms, source availability, and supported platforms vary by individual program. | Scriptable tools covering many established PET methods and data formats. | Heterogeneous interfaces, documentation, licences, and maintenance status; cite and version the exact program used. |
| **PMOD** | Long-standing commercial platform | Integrated image processing, VOI analysis, kinetic modelling, parametric mapping, and visualisation. | Proprietary commercial software. | Broad functionality, mature graphical interfaces, and common use across centres. | Source is not openly accessible; reproducibility depends on the licensed version, modules, and complete reporting of settings. |
| **COMKAT** | Muzic and Cornelius, 2001; Fang et al., 2010 | Compartmental modelling, simulation, parameter estimation, image display, ROI analysis, and model construction. | Historically distributed for non-commercial use; MATLAB-based. Current terms and availability should be verified. | Rich model specification with command-line and graphical interfaces; recognised historical role in PET/SPECT modelling. | Thielemans (2023) noted restricted contribution pathways and uncertain maintenance; redistribution and current support may be limited. |
| **DEPICT** | Gunn et al., 2002 | Data-driven estimation of regional parameters and parametric images using compartmental theory and basis-pursuit denoising. | Academic-use MATLAB distribution; registration and acceptance of restrictive terms required; redistribution and commercial use prohibited. | Seminal, theoretically documented approach that can estimate parameters and model order without prespecifying a compartmental structure. | Restricted, non-redistributable access and legacy MATLAB requirements limit transparent deployment and long-term reproducibility. |
| **Kinetic Imaging System (KIS)** | Legacy UCLA resource; included in Thielemans 2023 overview | Kinetic modelling and related analysis of dynamic PET/SPECT data in a dedicated Java-based environment. | Registration required and not redistributable, as reported by Thielemans (2023); current availability and exact terms could not be confirmed. | Historically provided a dedicated environment for quantitative kinetic analysis. | Access restrictions, Java-era deployment, and uncertain current status limit inspection, redistribution, and reproducible reuse. |
| **Tracer Kinetic Model Fitting (TKMF)** | Legacy UCLA web resource; included in Thielemans 2023 overview | Server-based fitting of regional PET/SPECT TACs using user-selected tracer kinetic models. | Password/registration-controlled server access; software not openly distributed. | Centralised execution reduced local installation requirements and supported multiple established models. | Restricted access, limited inspectability, and dependence on a remote service make exact versioning and long-term reproducibility difficult. |
| **MIAKAT** | Gunn et al., 2016; Searle et al., 2017 | Integrated PET workflow including motion correction, anatomical parcellation, blood/plasma input-function modelling, and regional or voxel-wise kinetic modelling. | MATLAB m-code historically distributed free for non-commercial use under a licence agreement; current public availability should be confirmed. | Broad quantitative workflow with graphical interfaces, standard workflows, and support for bespoke analyses. | Proprietary MATLAB and external dependencies, non-commercial terms, and uncertain current public distribution require exact version and licence reporting. |
| **Spamalize** | Legacy Waisman Laboratory software; included in Thielemans 2023 overview | SPECT, PET, and MRI display and analysis, with end-use applications and preprocessing/filtering utilities. | IDL-based; reported as non-commercial and not redistributable by Thielemans (2023). | Historically cross-platform and extensible through specialised in-house imaging utilities. | Legacy proprietary IDL dependency, restricted redistribution, and unclear maintenance; many components are not specific to kinetic modelling. |
| **Imager-4D** | Rowe et al., 2019 | Interactive visualisation and analysis of dynamic PET, including VOI extraction, Patlak analysis, and radiomic features. | Free downloadable Java/JavaFX application; no standard open-source licence was identified. | Cross-platform Java design, graphical dynamic-data exploration, and integrated quantitative and radiomic outputs. | Free availability is not equivalent to open source; source, redistribution terms, and automated workflow support are limited or unclear. |
| **imlook4d** | Long-standing MATLAB project; included in Thielemans 2023 overview | Four-dimensional PET/CT/MRI viewer with VOI analysis, pharmacokinetic modelling, format conversion, and drop-in MATLAB extensions. | Source publicly visible on GitHub; MATLAB required; no clear repository-level licence was identified. | Interactive 4D visualisation, broad format support, and straightforward extension through MATLAB scripts. | Public code without an explicit licence does not establish reuse or redistribution rights; GUI-driven workflows and MATLAB dependencies require careful documentation. |
| **FireVoxel** | Long-standing NYU software; project status updated 1 July 2026 | MRI, CT, PET, and SPECT display and analysis, including kinetic modelling, DICOM import, segmentation, ROI definition, and registration. | Non-commercial research-use software; Windows executable. Distribution, development, and support were suspended as of 1 July 2026. | Broad integrated image-analysis functionality with extensive documentation and tutorials. | Closed executable and non-commercial terms limit inspectability; the current project hold creates major availability and sustainability concerns. |
| **mfEVolve** | MultiFunctionalImaging research software; included in Thielemans 2023 overview | Multi-tracer dynamic PET processing, compartment-model fitting, tracer separation, and parametric imaging. | Research-use-only proprietary software; patent-protected methods and provider-controlled access. | Specialised support for simultaneous fitting of multiple tracers and rapid parametric modelling. | Not open source; access, licensing, implementation details, and independent version-specific validation must be documented. |
| **SAKE** | Veronese et al., 2013 | Stand-alone spectral analysis for kinetic estimation, dynamic PET quantification, and parametric analysis. | Official project page describes a free, licence-free distribution; Thielemans (2023) reported CC BY-NC-SA 3.0 with Windows/C++ and MATLAB Runtime. Formal current terms should be verified. | Dedicated implementation of established spectral-analysis methods with a stand-alone user interface. | Inconsistent licence descriptions, non-commercial restrictions reported in 2023, and platform/runtime constraints complicate reuse and redistribution. |
| **QModeling** | Lopez-Gonzalez et al., 2019 | SPM toolbox for SRTM, SRTM2, reference Patlak, and reference Logan analyses of dynamic PET. | GNU GPL v3 or later; MATLAB/SPM; registration required. Official page lists release 2.1. | Open-source copyleft licence, documented reference-region models, and graphical integration with SPM. | Registration gate and proprietary MATLAB dependency; tested MATLAB/SPM combinations are dated and the model set is focused on reference-region approaches. |
| **APPIAN** | Funck et al., 2018 | Automated PET preprocessing, quantification, and quality-control pipeline. | MIT; Python/Nipype; distributed through containers. | Modular automation, containers, documented pipeline execution, and extensibility. | Complex dependencies and external-tool integration require careful versioning and maintenance checks. |
| **Magia** | Karjalainen et al., 2020 | Automated brain PET processing using FreeSurfer/SPM, anatomical parcellation, Turku PET Centre modelling tools, metadata annotation, and quality control. | MIT-licensed source; MATLAB-based pipeline with multiple external dependencies. | Automated regional workflow, standardised parcellation and modelling, and published validation of automated reference-region generation. | Reproducibility depends on proprietary MATLAB and on exact versions/licences of FreeSurfer, SPM, and Turku tools; primarily designed for brain PET. |
| **NiftyPAD** | Jiao et al., 2023; highlighted by Thielemans 2023 | Dynamic PET kinetic analysis and parametric modelling in Python. | Apache License 2.0; Python. | Inspectable library code, reusable model implementations, and programmatic workflows. | Primarily a modelling library rather than a complete preprocessing-to-statistics pipeline. |
| **Dynamic PET** | Public Python project; included in Thielemans 2023 overview | Voxel-wise and regional analysis of reconstructed dynamic PET, including denoising, SUVR, Logan reference, and SRTM-family methods through a CLI and Python API. | MIT; Python 3.11+. | Scriptable CLI/API, public tests and documentation, and support for both image-level and regional workflows. | Model coverage is still expanding; several implementations are computationally intensive and the package is not a complete anatomical preprocessing pipeline. |
| **kinfitr** | Tjerkaski et al., 2020; Matheson et al., 2019 | Kinetic modelling of PET time-activity curves with diagnostics and outcome estimation. | MIT; R package. | Scriptable analyses, transparent model fitting, diagnostics, testing, and version control. | Focused on TAC/region-level modelling rather than full image preprocessing. |
| **bloodstream** | Public R/BIDS project; included in Thielemans 2023 overview | Configuration-driven processing of PET-BIDS blood data, including interpolation or model fitting and generation of input-function derivatives and reports. | MIT; R package; Docker and Apptainer deployment. | BIDS-aware inputs and outputs, automated reports, reusable configuration files, and containerised execution. | Limited to blood/input-function processing rather than tissue kinetic modelling or full image preprocessing; development versions should be pinned and cited. |
| **MITK-ModelFit** | Part of the MITK framework; included in Thielemans 2023 overview | General model fitting for dynamic imaging, including kinetic-model applications. | BSD-3-style open-source framework; C++ (verify the exact current module terms). | Embedded in a larger imaging toolkit with reusable infrastructure and a broader contributor base. | Broader than PET-specific modelling and may require substantial integration expertise. |
| **PETSurfer** | Greve et al., 2014, 2016 | Surface-based PET-MR registration, anatomical segmentation, partial-volume correction, and quantitative analysis. | Source available within the FreeSurfer distribution and its licence terms. | Tight integration with structural MRI and cortical-surface workflows. | Ecosystem-specific and not distributed under a standard permissive open-source licence. |
| **STARE** | Bartlett et al., 2022 | Blood-free estimation of net influx rate for irreversible PET tracers using Source-to-Target Automatic Rotating Estimation. | MATLAB code publicly available on GitHub; FSL/NIfTI dependencies; no clear repository-level licence was identified. | Published and validated method with a public, modular implementation that avoids arterial blood sampling. | Specialised to irreversible kinetics; licence is unclear and the released implementation was developed and validated with older MATLAB/FSL versions. |
| **PETPrep** | Emerging BIDS-oriented preprocessing projects | Automated PET preprocessing, registration, normalisation, ROI definition, and optional corrections. | Open/source-available and container-oriented; confirm the exact release and licence cited. | BIDS inputs, automation, containers, explicit provenance, and scalable execution. | Still maturing; feature, tracer, and validation coverage may differ across versions. |
| **PICNIC** | Emerging BIDS-oriented preprocessing projects | Automated PET preprocessing, registration, normalisation, ROI definition, and optional corrections. | Open/source-available and container-oriented; confirm the exact release and licence cited. | BIDS inputs, automation, containers, explicit provenance, and scalable execution. | Still maturing; feature, tracer, and validation coverage may differ across versions. |
| **PETFit** | Emerging BIDS App for kinetic modelling; manuscript/submission cited in the review article | Kinetic modelling of PET TAC and blood-input data with standardised inputs and outputs. | Open-source project; verify the exact public release, archived version, and licence before publication. | Direct BIDS integration, scriptable execution, containers, and version-controlled modelling. | Under active development; model-specific validation and a stable release citation are essential. |
